# PepBERT: Lightweight language models for bioactive peptide representation

**DOI:** 10.1101/2025.04.08.647838

**Authors:** Zhenjiao Du, Doina Caragea, Xiaolong Guo, Yonghui Li

**Author notes:** Corresponding author: Dr. Yonghui Li.

## Abstract

Protein language models (pLMs) have been widely adopted for various protein and peptide-related downstream tasks and demonstrated promising performance. However, short peptides are significantly underrepresented in commonly used pLM training datasets. For example, only 2.8% of sequences in the UniProt Reference Cluster (UniRef) contain fewer than 50 residues, which potentially limits the effectiveness of pLMs for peptide-specific applications. Here, we present PepBERT, a lightweight and efficient peptide language model specifically designed for encoding peptide sequences. Two versions of the model—PepBERT-large (4.9 million parameters) and PepBERT-small (1.86 million parameters)—were pretrained from scratch using four custom peptide datasets and evaluated on nine peptide-related downstream prediction tasks. Both PepBERT models achieved performance superior to or comparable to the benchmark model, ESM-2 with 7.5 million parameters, on 8 out of 9 datasets. Overall, PepBERT provides a compact yet effective solution for generating high-quality peptide representations for downstream applications. By enabling more accurate representation and prediction of bioactive peptides, PepBERT can accelerate the discovery of food-derived bioactive peptides with health-promoting properties, supporting the development of sustainable functional foods and value-added utilization of food processing by-products. The datasets, source codes, pretrained models, and tutorials for the usage of PepBERT are available at https://github.com/dzjxzyd/PepBERT.

## 1. Introduction

Bioactive peptides—protein fragments that exhibit beneficial biological activities such as antioxidant, anticancer, and antimicrobial functions—are of considerable interest in nutraceutical, therapeutic, and industrial applications (Du et al., 2023a; Du and Li, 2022). Traditional wet-lab methods for peptide discovery are labor-intensive, expensive, and time-consuming; thus, machine learning approaches have become popular for virtual screening at a large scale (Huang et al., 2023). A key step in developing effective machine learning models for peptide discovery is obtaining robust numerical representations of peptides (Du et al., 2023a).

With the advent of transformer architectures, large language models (LLMs) have achieved remarkable success in text representation by leveraging self-supervised learning on massive datasets (Zhang et al., 2024). Biological sequences (e.g., DNA, RNA, proteins) share a similar sequential structure with natural language, making them amenable to such techniques (Zhang et al., 2024). Protein language model (pLM) is a type of LLM trained on millions of protein sequences, and has been widely applied to a variety of prediction tasks, including structure prediction (Lin et al., 2023), solubility (Thumuluri et al., 2022), function annotation (Xiang et al., 2024), etc. Peptides, which are short fragments of proteins, have also benefited from the growing adoption of pLMs (e.g., ESM-2 (Lin et al., 2023)). However, the percentage of peptide sequences in the training dataset for pLM remains scarce. For example, in widely used dataset UniRef100 (Suzek et al., 2014), only 0.06% of sequences are shorter than 50 residues, which limits their effectiveness for peptide-specific tasks. Moreover, employing a full-scale protein language model for peptides may be unnecessarily complex and computationally intensive given the shorter sequence lengths involved.

Pretraining a peptide language model on a peptide dataset has the potential to enhance the representation of peptides and further advance the peptide prediction model development. Previous work, PeptideBERT (Guntuboina et al., 2023), has contributed to peptide language model development. However, it focused on fine-tuning the pretrained ProteinBERT (Brandes et al., 2022) for individual downstream tasks, rather than training a peptide-specific language model from scratch or fine-tuning a general-purpose peptide language model to enable broader representation learning across the peptide discovery community. To date, there is no dedicated pretrained peptide language model specifically for embeddings of short amino acid sequences. To meet this demand, we developed PepBERT—a lightweight peptide language model tailored specifically for peptide representation.

In this study, we adopted a bidirectional encoder representation from transformers (BERT)-based architecture with a masked language modeling strategy and pretrained our models with two variants (PepBERT-small with 1.86 million parameters and PepBERT-large with 4.9 million parameters) on four datasets comprising between 1.97 and 19.2 million peptide sequences. The pretrained PepBERT models were released at Hugging Face (https://huggingface.co/dzjxzyd) with detailed tutorials for usage for peptide embedding. We then evaluated their performance on nine bioactive peptide datasets via transfer learning. Compared with the smallest variant of ESM-2 (7.5 million parameters), our models demonstrate equal or superior performance on 8 out of 9 datasets. PepBERT offers an alternative to pLMs for peptide representation, aiming to further enhance the performance of peptide prediction models. Its lightweight nature makes it well-suited for broader applications with reduced computational cost.

## 2. Methods

### 2.1 Datasets

To train a peptide language model, we first built peptide datasets excluding those long protein sequences. We curated our datasets based on four publicly available protein datasets: UniProt Archive (UniParc), UniRef 100, UniRef90, and UniRef50. UniRef database is the clustered sets of sequences from UniProt Knowledgebase (UniProtKB), and UniRef100, UniRef90 and UniRef50 are filtered at varying sequence identity thresholds (100%, 90%, 50%). In contrast, UniParc is a comprehensive archive of protein sequences that acts as a historical record (Suzek et al., 2014). Only sequences with lengths of 50 residues or fewer were retained. As a result, four datasets were obtained (Table 1), each representing peptide sequences at different scales. These datasets were subsequently compared to evaluate their impact on model performance. The detailed length distributions of the sequences in the collected datasets are illustrated in Figure 1.

**Table 1.** Four datasets for PepBERT pretraining

| Datasets | Sequence length | Number of sequences (M) | Number of amino acids (M) | Download links |
| --- | --- | --- | --- | --- |
| UniParc | 2-50 | 19.2 | 78 | https://huggingface.co/datasets/dzjxzyd/UniParc_len_0_50 |
| UniRef100 | 2-50 | 2.7 | 94 | https://huggingface.co/datasets/dzjxzyd/UniRef100_len_0_50 |
| UniRef90 | 2-50 | 2.35 | 108 | https://huggingface.co/datasets/dzjxzyd/UniRef90_len_0_50 |
| UniRef50 | 2-50 | 1.97 | 663 | https://huggingface.co/datasets/dzjxzyd/UniRef90_len_0_50 |

**Figure 1.**
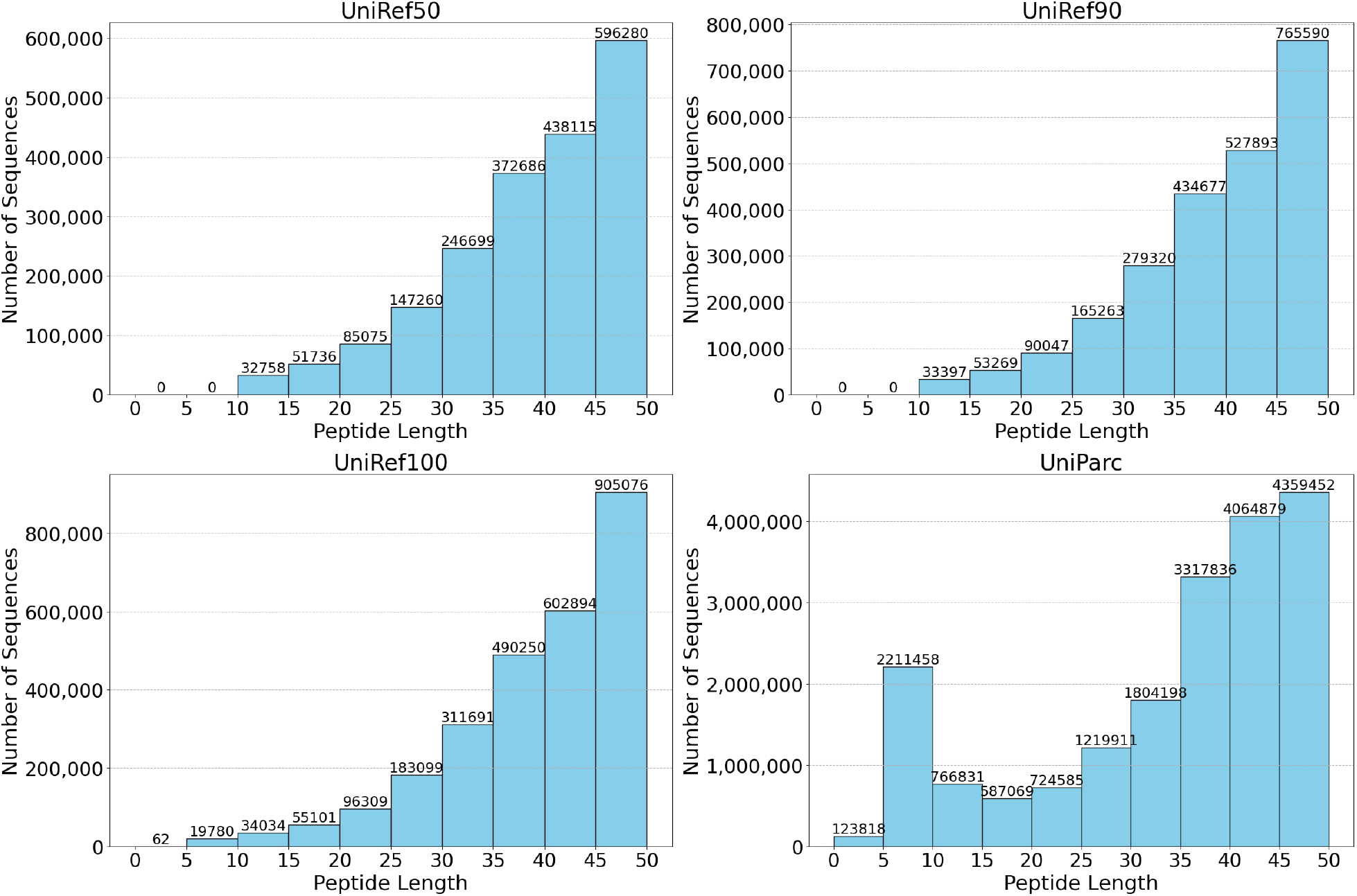
Peptide length distribution in each of the four datasets used for pretraining

### 2.2 Model architecture and pretraining setup

With the peptide specific dataset, we chose a similar model architecture as that of ESM-2 model with a few modifications. Since all peptides are 50 residues or shorter, we directly employed sinusoidal positional embeddings. Two architectures were developed: PepBERT-small with 1.86 million parameters and 160-dimensional embeddings and PepBERT-large with 4.94 million parameters and 320-dimensional embeddings. Both models consist of 8 encoder layers, each featuring an 8-head multi-head self-attention mechanism. The feed-forward network in each layer uses an intermediate dimension of 640 units.

For optimization, we used the AdamW optimizer with hyperparameters β_1_ = 0.9, β_2_ = 0.98, ε = 10^−8^, and a weight decay of 0.01. The learning rate was warmed up over the first 2000 steps to a peak of 4×10^−4^ and then decayed linearly to one-tenth of the peak value over 90% of the training duration. The maximum input length was fixed at 50 amino acids. During tokenization, special tokens were incorporated to denote specific positions or purposes in the sequence: [SOS] for the start of a sequence, [EOS] for the end, [PAD] for padding, [UNK] for unknown tokens, and [MASK] for masked positions. For the masked language modeling task, 15% of amino acids in each peptide were randomly replaced by the [MASK] token, and cross-entropy loss was computed over the predictions.

A validation set comprising 0.1% of the sequences was held out, and model checkpoints were saved after each epoch. Specifically, there were 19180, 2698, 2349, and 1970 peptide sequences in the validation sets of UniParc, UniRef 100, UniRef90, and UniRef50, respectively. The checkpoint with the lowest validation loss over 20 epochs was selected as the final model for each architecture and dataset. The training scripts are available at https://github.com/dzjxzyd/PepBERT for the PepBERT-small and PepBERT-large models, respectively. Corresponding to the 4 datasets used for training, 4 PepBERT-small models and 4 PepBERT-large models were pretrained, for a total of eight pretrained models that can be downloaded from Hugging Face at https://huggingface.co/dzjxzyd, along with a tutorial for generating peptide embeddings.

### 2.3 Transfer learning on bioactive peptide datasets and performance comparison

Nine bioactive peptide classification datasets (Table S1) were selected from previous studies in our laboratory (Du et al., 2023b; Kumar et al., 2025), restricted to sequences shorter than 50 residues. The pretrained PepBERT models were used to generate peptide embeddings without finetuning for different downstream tasks. For benchmarking, we compared with the smallest variant of ESM-2—a well-established protein language model with 7.5 million parameters and 320-dimensional output embeddings (Lin et al., 2023). Each dataset was randomly shuffled and split into training and test sets (80:20 ratio), repeated ten times. To ensure a fair comparison of the embeddings and to minimize the influence of the classifier itself, logistic regression (max_iter=1000) was used for classification model development. Statistical comparisons between ESM-2 and PepBERT performances were conducted using a t-test (p < 0.05), with evaluation metrics including accuracy (ACC), balanced accuracy (BACC), sensitivity (Sn), specificity (Sp), Matthews correlation coefficient (MCC), and area under the curve (AUC).

## 3. Results and discussion

In Figure 2, we can clearly observe the constant improvements to the validation loss with the increased dataset size for both models. Specifically, the PepBERT trained on UniParc with 19.2 million peptide sequences had the lowest loss in PepBERT-small and PepBERT-large, respectively. A similar observation was also reported in other large language model development studies (Ahmad et al., 2022; Moret et al., 2023). Based on Chinchilla scaling laws: 20 tokens-per-parameter ratios as the optimal for foundation model development, PepBERT-large and PepBERT-small needed 98 million and 37.2 million tokens, respectively, which was fulfilled in most cases in our study (Table 1). We believe that the improvements can be mostly attributed to the increasing diversity of the dataset. Larger datasets, particularly the UniParc dataset (Figure 1), preserved more peptides with only a few residues (Figure 1). Additionally, PepBERT-large, which has more learnable parameters, outperformed PepBERT-small. The results are consistent with the previous studies in pLMs, like ProtTrans (Elnaggar et al., 2021) and ESM (Lin et al., 2023), where larger datasets and larger models generally lead to better performance. This aligns with the well-established scaling laws observed in natural language processing (Kaplan et al., 2020). This also suggests that the learning capacity of PepBERT-small may be insufficient to capture the patterns in such large datasets.

**Figure 2.**
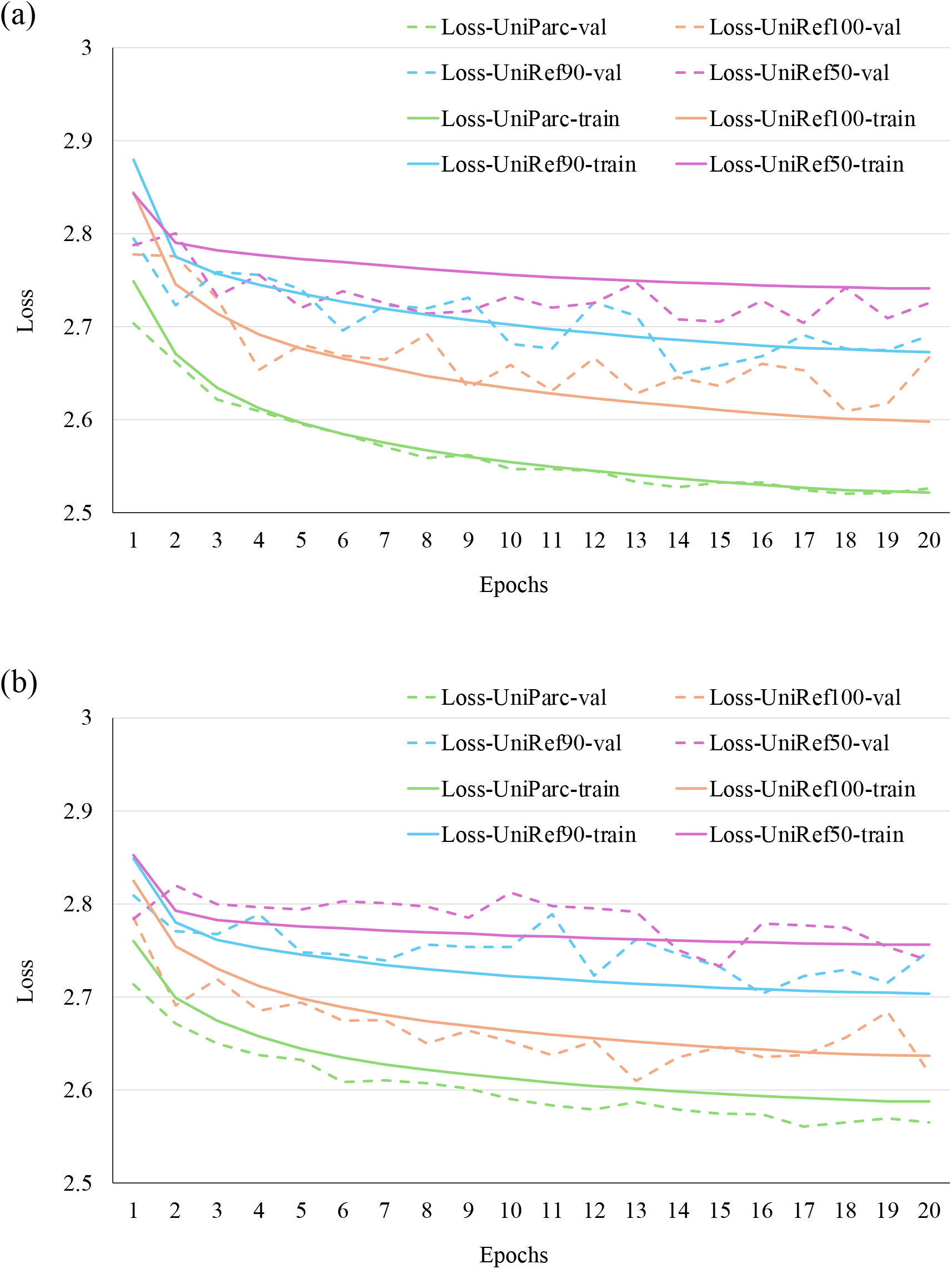
Validation loss curves of the PepBERT-small (a) and PepBERT-large (b) models.

The checkpoints corresponding to the lowest validation loss were then used for downstream transfer learning tasks, alongside the benchmark ESM-2 model. As shown in Tables 2 and 3, the models pretrained on UniParc achieved the best overall performance among the models pretrained on our four datasets. Specifically, compared with ESM-2, PepBERT-large-UniParc outperformed it in 3 of 9 datasets (p < 0.05) and reached equivalent performance in 5 of 9 datasets with no significant differences in balanced accuracy (BACC). In contrast, PepBERT-small-UniParc demonstrated significantly better performance in 1 of 9 datasets (p < 0.05) and achieved comparable performance in 7 of 9 datasets. Detailed performance metrics are provided in Table S2.

**Table 2.** Comparison of PepBERT-small with ESM-2 on 9 bioactive peptide classification tasks.

| Prediction tasks | ESM-2 | PepBERT-small-UniParc | PepBERT-small-UniRef100 | PepBERT-small-UniRef90 | PepBERT-small-UniRef50 |
| --- | --- | --- | --- | --- | --- |
| ACE inhibitory peptide | 0.756±0.018 | 0.750±0.018** | 0.747±0.020** | 0.757±0.014** | 0.755±0.021** |
| Quorum sensing peptide | 0.881±0.044 | 0.890±0.031** | 0.893±0.022** | 0.885±0.028** | 0.874±0.051** |
| Anticancer peptide-main | 0.926±0.011 | 0.916±0.010** | 0.892±0.014* | 0.896±0.013* | 0.901±0.012* |
| Anticancer peptide-alternative | 0.690±0.027 | 0.720±0.025*** | 0.708±0.015** | 0.699±0.019** | 0.699±0.026** |
| Tumor T cell antigens | 0.634±0.034 | 0.658±0.026** | 0.656±0.032** | 0.675±0.027*** | 0.650±0.028** |
| Blood-brain barrier penetrating peptides | 0.750±0.050 | 0.715±0.059** | 0.692±0.060* | 0.746±0.081** | 0.740±0.037** |
| Neuropeptide | 0.848±0.015 | 0.837±0.017** | 0.831±0.011* | 0.822±0.015* | 0.831±0.014* |
| Toxicity | 0.883±0.014 | 0.883±0.008** | 0.877±0.010** | 0.874±0.008** | 0.883±0.011** |
| Cell penetrating peptide | 0.855±0.007 | 0.836±0.010* | 0.836±0.010* | 0.831±0.009* | 0.837±0.007* |

Note: Performance comparison is based on balanced accuracy; \*\*\*: the performance of this model at the corresponding dataset was significantly higher than that of ESM-2 model ( $p<0.05$ ); \*\*: the performance of this model at the corresponding dataset had no difference with that of ESM-2 model ( $p<0.05$ ); \*: the performance of this model at the corresponding dataset was significantly lower than that of ESM-2 model ( $p<0.05$ ).

**Table 3.**
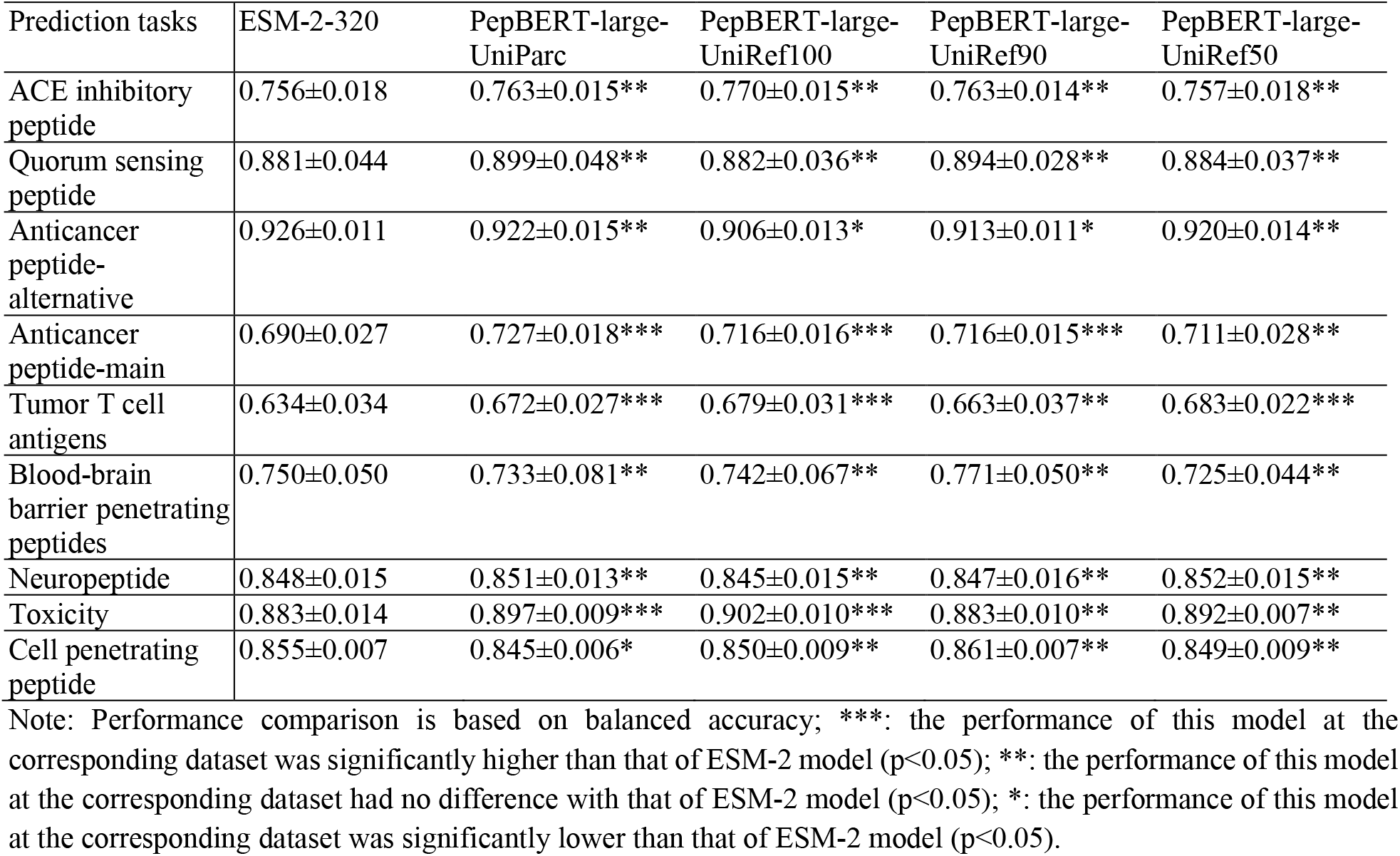
Comparison of PepBERT-large with ESM-2 on 9 bioactive peptide classification tasks.

| Prediction tasks | ESM-2-320 | PepBERT-large-UniParc | PepBERT-large-UniRef100 | PepBERT-large-UniRef90 | PepBERT-large-UniRef50 |
| --- | --- | --- | --- | --- | --- |
| ACE inhibitory peptide | 0.756±0.018 | 0.763±0.015** | 0.770±0.015** | 0.763±0.014** | 0.757±0.018** |
| Quorum sensing peptide | 0.881±0.044 | 0.899±0.048** | 0.882±0.036** | 0.894±0.028** | 0.884±0.037** |
| Anticancer peptide-alternative | 0.926±0.011 | 0.922±0.015** | 0.906±0.013* | 0.913±0.011* | 0.920±0.014** |
| Anticancer peptide-main | 0.690±0.027 | 0.727±0.018*** | 0.716±0.016*** | 0.716±0.015*** | 0.711±0.028** |
| Tumor T cell antigens | 0.634±0.034 | 0.672±0.027*** | 0.679±0.031*** | 0.663±0.037** | 0.683±0.022*** |
| Blood-brain barrier penetrating peptides | 0.750±0.050 | 0.733±0.081** | 0.742±0.067** | 0.771±0.050** | 0.725±0.044** |
| Neuropeptide | 0.848±0.015 | 0.851±0.013** | 0.845±0.015** | 0.847±0.016** | 0.852±0.015** |
| Toxicity | 0.883±0.014 | 0.897±0.009*** | 0.902±0.010*** | 0.883±0.010** | 0.892±0.007** |
| Cell penetrating peptide | 0.855±0.007 | 0.845±0.006* | 0.850±0.009** | 0.861±0.007** | 0.849±0.009** |

Note: Performance comparison is based on balanced accuracy; \*\*\*: the performance of this model at the corresponding dataset was significantly higher than that of ESM-2 model ( $p<0.05$ ); \*\*: the performance of this model at the corresponding dataset had no difference with that of ESM-2 model ( $p<0.05$ ); \*: the performance of this model at the corresponding dataset was significantly lower than that of ESM-2 model ( $p<0.05$ ).

It is worth noting that while PepBERT is designed to generate improved representations for short amino acid sequences (i.e., peptides), there are still underrepresented groups, particularly peptides with 2 to 5 residues. These ultrashort peptides are especially important in peptide-based small molecule drug development (Du and Li, 2022). However, training an LLM using masked language modeling or next-token prediction is inherently challenging on those extremely short sequences, considering the mechanisms behind LLMs. There is less contextual information for those short peptides, and hence, the quality of representations generated by LLM for these peptides may be limited, and expectations should be moderated accordingly. One potential solution is to adopt alternative linear notations—such as the simplified molecular-input line-entry system (SMILES) or self-referencing embedded strings (SELFIES) (Krenn et al., 2020) —to transform shorter amino acid sequences into longer, text-like formats. These alternative notations also facilitate the extension to modified or cyclic peptides. Wang et al. recently introduced PepDoRA, a peptide representation model based on SMILES notation (Wang et al., 2024). PepDoRA fine-tunes ChemBERTa on 100,000 peptide sequences using weight-decomposed low-rank adaptation (DoRA) using a masked language model objective to generate optimized embeddings for downstream tasks involving both modified and unmodified peptides. Interestingly, when applied solely to unmodified peptides, PepDoRA did not show significant performance improvements over PeptideBERT, highlighting both the potential and limitations of alternative notational approaches (Wang et al., 2024).

## 4. Conclusion

In summary, we present a lightweight peptide language model, PepBERT, with two versions. Both models achieved promising performance across a range of downstream bioactive peptide prediction tasks, establishing their potential as efficient tools for peptide representation and providing a foundation for more efficient discovery and application of bioactive peptides in food systems. The datasets, source codes, pretrained models, and tutorials for usage of PepBERT are available at https://github.com/dzjxzyd/PepBERT. Nonetheless, more robust optimization, particularly through hyperparameter tuning, could further enhance their effectiveness. Exploring alternative notation systems for peptide language model development, such as SMILES or SELFIES, can be a worthy direction for future research. One challenge arises from the conserved structural regularity of peptide backbones. While this invariance simplifies the pattern recognition in canonical SMILES encodings, it can also introduce redundant token patterns that undermine the effectiveness of transformer-based attention mechanisms. To overcome this, a critical future direction involves designing adaptive tokenization schemes that dynamically modulate the granularity of sequential information encoding to maintain balance between sparse biological semantics (as in amino acid sequences) and chemically dense but repetitive notations like SMILES.

## Acknowledgements

This is contribution No. 26-004-J from the Kansas Agricultural Experimental Station. This work was supported in part by the National Science Foundation (2419880). Research reported in this publication was also partially supported by the Cognitive and Neurobiological Approaches to Plasticity Center of Biomedical Research Excellence of the NIH under grant No. P20GM113109. The content is solely the responsibility of the authors and does not necessarily represent the official views of the NIH.

## Data availability

Data will be made available on request.

